# Technology-enhanced learning in undergraduate neuroscience education: tractography-based virtual dissection in psychology

**DOI:** 10.64898/2026.09.21.753122

**Authors:** Diana López-Barroso, María José Torres-Prioris

## Abstract

**Background:** Neuroanatomy poses a significant challenge for Psychology students due to its spatial and conceptual complexity. Educational approaches that enhance the relevance and visualization of neuroanatomical content may improve students’ learning experiences. This study implemented a tractography-based activity focused on the virtual dissection of the arcuate fasciculus, a major white matter pathway, in undergraduate Psychology students and examined the relationships between perceived learning and students’ perceptions of utility, difficulty and handling, and organizational aspects of the activity.

**Methods:** First-year undergraduate Psychology students participated in a two-session tractography-based activity combining instruction on white matter anatomy and diffusion tractography with a hands-on virtual dissection of the arcuate fasciculus using research-grade software routinely employed in neuroscience research. Following the activity, students completed an anonymous questionnaire assessing perceived learning, utility, difficulty and handling, and organizational aspects of the activity. Pearson correlations, multiple regression analyses, and relative importance analyses were performed.

**Results:** Sixty-eight students completed the questionnaire. Students reported generally positive perceptions of the activity across the evaluated dimensions, with perceived learning receiving the highest mean score (M = 3.44, SD = .85). Perceived utility showed the strongest association with perceived learning (r = .64, p < .001). The regression model explained 41% of the variance in perceived learning (R² = .41, adjusted R² = .38, p < .001). Perceived utility was the only significant predictor in the model (β = .59, p = .001), accounting for 67.9% of the model’s explained variance.

**Conclusions:** The findings support the feasibility of integrating authentic neuroimaging tools into undergraduate neuroanatomy teaching. Students who perceived the activity as more useful also reported higher perceived learning outcomes, with perceived utility emerging as the strongest predictor of perceived learning. In contrast, perceived difficulty and handling, and organizational aspects did not make significant independent contributions. These results suggest that students’ perceptions of educational relevance may play an important role in technology-enhanced STEM learning experiences.

## 1. Introduction

Courses within the fields of Psychobiology and Neuroscience represent a core component in undergraduate Psychology education, as they provide the biological foundations necessary for understanding human behavior, cognition, and mental processes. However, these courses, particularly those with a strong neuroanatomical component, are often perceived as challenging by students (Edwards-Bailey et al., 2022) due to the inherent complexity of brain structure and organization, especially during the early years of the degree. These challenges may be further intensified when students do not readily perceive the value of neuroanatomical knowledge within their education. In a recent study, 64% of second-year Psychology students reported *neurophobia* (Garcés-Arilla et al., 2025), defined as apprehension toward neuroscience-related subjects (Jozefowicz, 1994). To address this challenge, educational approaches should facilitate the understanding of complex neuroanatomical structures and their spatial relationships. In this context, teaching strategies that support the visualization of brain anatomy and connectivity may help students overcome some of the difficulties traditionally associated with learning neuroanatomy.

A central challenge in neuroanatomy education is helping students develop accurate three-dimensional representations of the anatomical complexity of the brain. More broadly, difficulties in understanding complex spatial structures are common across STEM disciplines and have motivated the development of technology-enhanced learning environments that support visualization and active exploration of abstract concepts. Traditional approaches to neuroanatomy teaching have largely relied on two-dimensional representations found in atlases and textbooks (Figure 1A), requiring students to mentally reconstruct three-dimensional (3D) anatomical relationships from separate views. More recently, physical 3D plastic models have been incorporated into anatomy teaching to facilitate the visualization and manipulation of major neuroanatomical structures (Figure 1B). However, these models usually provide only simplified representations of brain anatomy, with limited anatomical resolution and anatomical variability, as well as a reduced capacity to depict the complexity of white matter organization. In some cases, educational models may even promote oversimplified or partially inaccurate spatial representations of certain structures^1^. These limitations, together with the spatial complexity of neuroanatomical systems, highlight the potential value of technology-enhanced approaches that provide interactive representations (Čavka M et al., 2025; Yammine & Violato, 2015). Among these approaches, virtual tractography enables the three-dimensional visualization and dynamic exploration of white matter pathways reconstructed from diffusion-weighted magnetic resonance imaging (DTI-MRI) data, providing a direct representation of the fiber bundle trajectories and their spatial organization (Figure 1C). Beyond visualization, tractography-based activities require students to identify anatomical landmarks, define regions of interest (ROIs), reconstruct fiber pathways, and evaluate the anatomical plausibility of the resulting tract reconstructions. Such tasks engage spatial reasoning and scientific data interpretation skills that are characteristic of authentic STEM learning environments. Consistent with this potential, previous research suggests that three-dimensional visualization can support anatomical learning and student engagement (Chystaya et al., 2022) and that interactive virtual 3D tools may improve learning outcomes for complex spatial topics compared with traditional instructional approaches (Elsayed et al., 2025). In addition, because tractography is routinely used in contemporary neuroscience research, it provides students with exposure to an authentic scientific tool that may help bridge the gap between classroom learning and real-world scientific practice.

**Figure 1.**
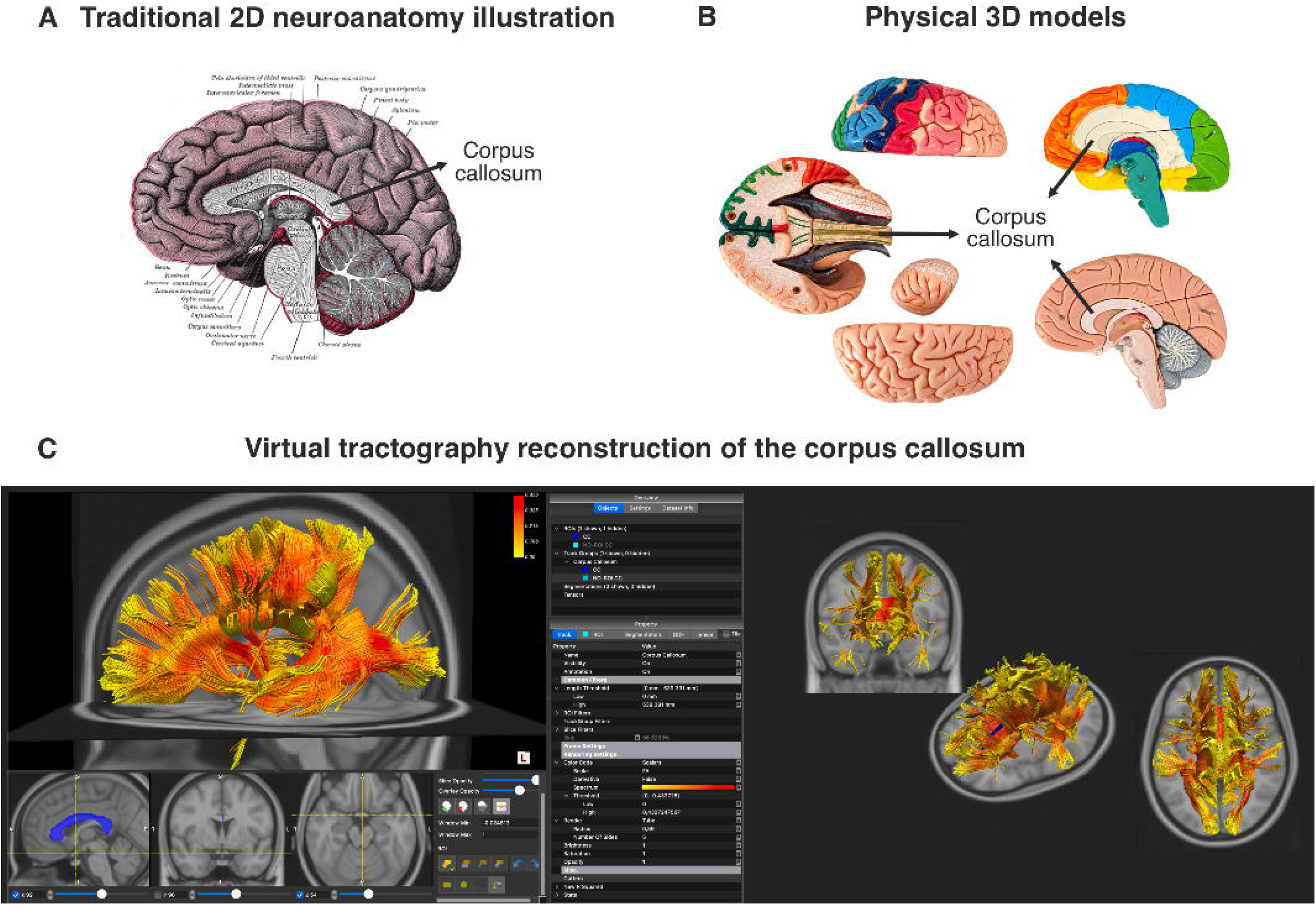
Three-panel comparison of educational resources for learning neuroanatomy and white matter anatomy, illustrating the progression from two-dimensional neuroanatomical representations to three-dimensional virtual tractography. **(A)** Traditional two-dimensional illustration of midsagittal brain section of the human brain, highlighting the corpus callosum (arrow). Such illustrations require learners to mentally reconstruct three-dimensional (3D) anatomical representations from planar images. **(B)** Example of physical 3D brain plastic models. Although it facilitates spatial understanding through direct manipulation and visualization of major brain structures, it provides only a simplified representation of white matter pathways and certain complex neuroanatomical structures. For instance, the corpus callosum (arrows) is represented in a non-intuitive manner, making its anatomical course and spatial trajectory difficult to appreciate. **(C)** Virtual tractography dissection of the corpus callosum using specialized software. This approach enables interactive 3D exploration of brain anatomy and white matter organization, including virtual dissection, rotation, zooming, and selective visualization of specific pathways, thereby facilitating the development of accurate mental representations of brain organization. Source for Panel A: Henry Vandyke Carter, illustration in Gray H. Anatomy of the Human Body, Plate 720 (1918). Public domain.

In line with these considerations, the present study describes and evaluates a technology-enhanced learning activity implemented in a first-year Psychobiology course within an undergraduate Psychology program. The activity integrated instruction on white matter anatomy and diffusion MRI tractography with hands-on virtual dissection of the arcuate fasciculus using research-grade neuroimaging software routinely employed in neuroscience research. The study had two main aims: (1) to characterize students’ perceptions of a technology-enhanced learning activity involving an authentic scientific tool, including perceived learning, utility, difficulty and handling, and organizational aspects; and (2) to identify which of these dimensions were most strongly associated with perceived learning outcomes. By addressing these aims, the study contributes to the growing literature on technology-enhanced and visualization-based learning in STEM education while providing evidence regarding the educational use of authentic neuroimaging tools in undergraduate teaching.

## 2. Material and methods

### 2.1. Educational context

The study was conducted within the course *Fundamentals of Psychobiology*, a compulsory first-year course in the undergraduate Psychology program at the University of Malaga. This course introduces students to the basic neurobiological foundations of behavior, including brain structure, functional organization, and fundamental neuroanatomical concepts. Participants were first-year undergraduate Psychology students (n = 68), with no prior exposure to formal neuroanatomy instruction. The teaching innovation was implemented as part of the practical component of the course, with the aim of complementing traditional lecture-based teaching with hands-on, experiential, and technology-enhanced learning. The activity was designed to promote spatial understanding of white matter anatomy while engaging students with an authentic neuroimaging tool commonly used in neuroscience research. Following completion of the activity, students were invited to complete a brief anonymous questionnaire designed to collect feedback on the educational experience and inform future improvements to the activity. Participation was voluntary, and no personally identifying information was collected. Ethical approval was obtained from the Institutional Review Board of the University of Malaga (Reference: 233-2026-H).

### 2.2. Procedure

The teaching activity was implemented in two sequential sessions. In the first session, the instructor introduced the activity’s objectives and provided the theoretical background required for it. This included an overview of neuroimaging techniques, with emphasis on DTI-MRI and the principles of tractography. The main categories of white matter tracts (commissural, association, and projection fibers) were also introduced, together with their principal anatomical connections and functional significance. In addition, the instructor, who has previous experience in DTI-MRI and tractography research, conducted a live demonstration of a virtual white matter dissection, illustrating the loading of a tractography dataset and the basic functionalities of the software. This demonstration aimed to familiarize students with the software environment and navigation tools before the practical session, as visualizing and manipulating tractography data can be highly abstract for individuals with no prior experience in neuroimaging or 3D anatomical representations. The virtual dissection was performed with TrackVis software (Wang et al., 2007). TrackVis is a freely available research-grade software package used for the visualization, dissection, and analysis of DTI-MRI tractography data in cognitive and clinical neuroscience research. TrackVis was selected because it is freely available, supports ROI-based tractography visualization and virtual dissection, and provides a relatively accessible interface for introducing students to tractography workflows. By allowing users to interactively reconstruct, explore, and virtually dissect white matter pathways, it provides access to analyses and visualizations typically performed in contemporary neuroscience research environments.

The second session was conducted in a computer laboratory and involved a hands-on practical activity. Due to time constraints inherent to the course structure, the activity focused on a single, functionally meaningful white matter tract as a proof of concept, the arcuate fasciculus. This white matter tract was selected due to its well-characterized anatomical trajectory and its established role in language processing (López-Barroso et al., 2013). To facilitate learning and reduce the initial technical demands associated with the software, students were provided in advance with a written step-by-step tutorial covering both the use of the TrackVis software and the anatomical procedures required for the dissection of the arcuate fasciculus, as well as a video tutorial specifically developed for the activity. These materials included detailed instructions for TrackVis navigation, anatomical localization of the Broca, Wernicke, and Geschwind ROIs, ROI-combination procedures, streamline cleaning using exclusion ROIs, and reconstruction of the different segments of the arcuate fasciculus. Both the written guide and the video tutorial are publicly available at https://doi.org/10.5281/zenodo.22129828, enabling interested educators and researchers to access the instructional materials used during the implementation of the activity. Their availability supports the reproducibility and potential transferability of the educational activity to other undergraduate teaching contexts. A practice tractography dataset was also made available before the class session, allowing students to familiarize themselves with the software environment if desired. During the practical session, all students worked with a second preprocessed whole-brain tractography dataset provided by the instructor.

### 2.3. Virtual dissection protocol

The activity was based on the virtual dissection of the arcuate fasciculus using a three-ROI protocol according to the model proposed by Catani and collaborators (Catani et al., 2005; López-Barroso et al., 2013). This model distinguishes a direct long segment and two indirect segments connecting frontal, parietal, and temporal language-related regions and has been widely used to characterize the anatomical organization of the arcuate fasciculus.

Students first loaded a whole-brain tractogram (Figure 2A) together with anatomical reference images (T1-weighted, fractional anisotropy [FA], and color-coded FA maps) (Figure 2B). Based on the provided anatomical landmarks, three cortical regions corresponding to classical language-related areas were identified in the FA color map and with the help of the T1-weighted image: Broca’s territory, which encompassed the inferior frontal gyrus and the ventral portion of the premotor cortex; Wernicke territory, which included the middle and posterior portions of the superior and middle temporal gyri; and Geschwind territory, which comprised the angular and supramarginal gyri (see Figure S1). Three ROIs were subsequently drawn, corresponding to each territory: the Broca ROI, the Wernicke ROI, and the Geschwind ROI (Figure 2C), and three tracts were generated, one from each ROI (Figure 2D). Then, by combining each pair of ROIs, students reconstructed the long direct segment, the posterior indirect segment, and the anterior indirect segment (Figure 2E). When necessary, exclusion ROIs (No-ROIs) were added to remove anatomically implausible streamlines (Figure 2F). The resulting tract reconstructions were visually inspected and manipulated in 3D space using axial, coronal, sagittal, and 3D views, as illustrated in Figure 1C using the corpus callosum as an example. Finally, students customized the rendering of the reconstructed tracts by assigning different colors to each segment and rendering streamlines as tubes, thereby facilitating the visual distinction of the three segments and enhancing the three-dimensional appreciation of their trajectories (Figure 2G). Due to time constraints within the course schedule, the virtual dissection was limited to the left hemisphere.

**Figure 2.**
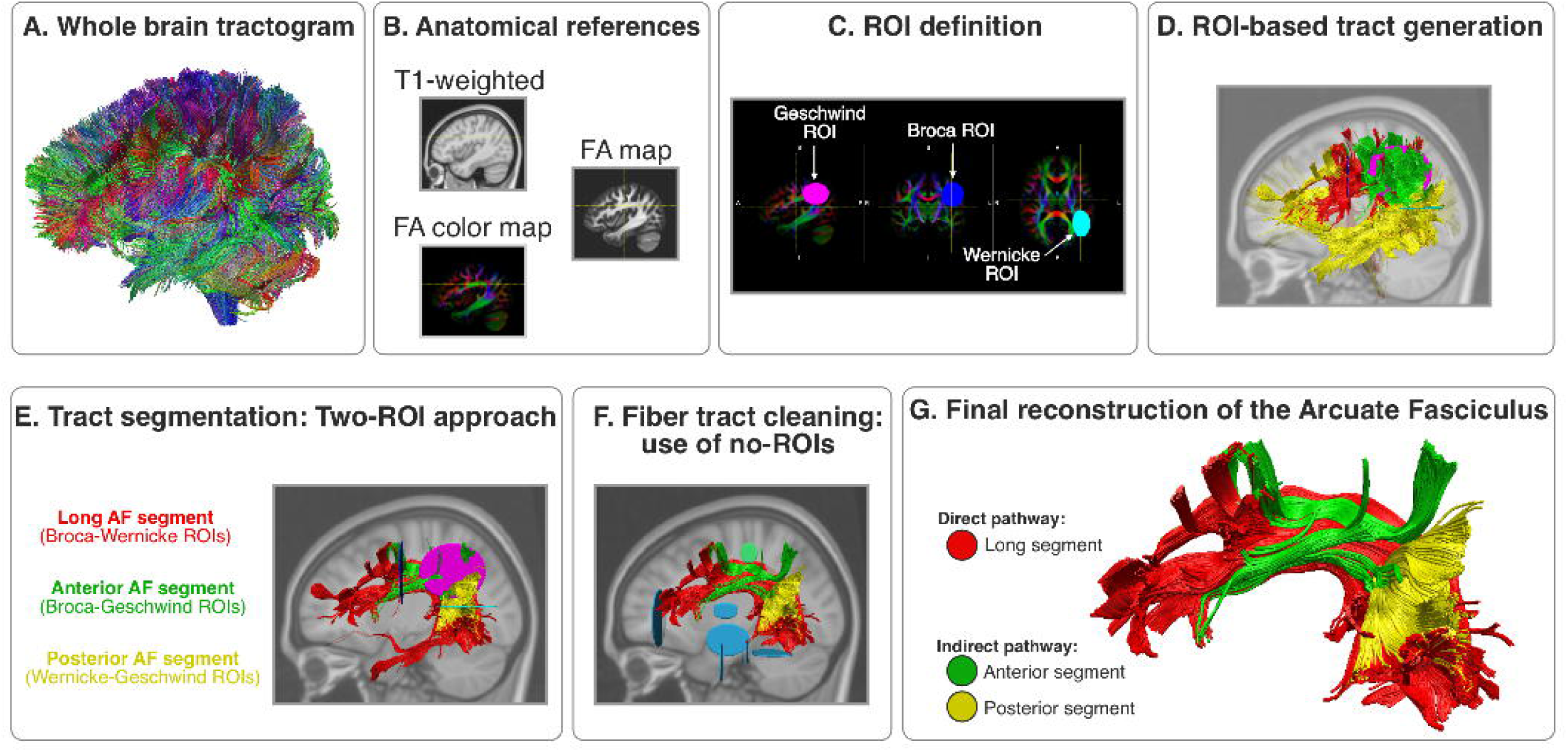
Virtual dissection workflow of the left arcuate fasciculus using TrackVis from a preprocessed tractogram. **(A)** Whole-brain tractogram used as the starting dataset for the practical activity. **(B)** Anatomical reference images, including T1-weighted, fractional anisotropy (FA), and color-coded FA maps, used to guide ROI placement **(C)**. Students identified the Broca, Geschwind, and Wernicke territories by following anatomical landmarks across axial, coronal, and sagittal views. **(D)** Initial ROI-based tract generation. For each ROI, streamlines intersecting the selected region were extracted from the whole-brain tractogram and visualized using different colors. **(E)** Reconstruction of the arcuate fasciculus segments using a two-ROI approach: the long direct segment connecting Broca’s and Wernicke’s territories (red), the anterior indirect segment connecting Broca’s and Geschwind’s territories (green), and the posterior indirect segment connecting Geschwind’s and Wernicke’s territories (yellow). **(F)** Tract refinement through the use of exclusion ROIs (No-ROIs) to remove anatomically implausible streamlines. **(G)** Final reconstruction of the left arcuate fasciculus showing the direct and indirect pathways after segmentation and cleaning. Streamlines are displayed as tubes to facilitate visualization of the tract’s 3D organization.

Throughout the activity, students directly manipulated tractography data, identified anatomical landmarks, created and edited ROIs, isolated white matter pathways, and explored their anatomical trajectories in 3D. These tasks required students to combine spatial reasoning, anatomical knowledge, and data interpretation skills during the reconstruction process. The practical session focused on the virtual dissection of the arcuate fasciculus using tractography data, rather than on its clinical relevance or cognitive functions, which had already been covered in previous theoretical lectures. The educational objectives were to develop students’ understanding of white matter anatomy and the three-dimensional organization of fiber pathways, while providing hands-on experience with ROI-based virtual tract reconstruction. Students submitted a report that included screenshots of their dissections and answers to conceptual questions about the anatomy of the arcuate fasciculus. The instructor provided guidance throughout the activity, and students were allowed to consult their class notes and ask questions. Thus, the assessment was not a formal examination but focused primarily on the practical dissection activity and students’ understanding of the arcuate fasciculus.

### 2.4. Evaluation of the activity

Upon completion of the practical activity, students completed an anonymous Learning Experience Questionnaire. The instrument consisted of 15 items rated on a 5-point Likert scale (1 = strongly disagree, 5 = strongly agree). The questionnaire was specifically developed to assess students’ perceptions of four dimensions of the educational experience: perceived learning outcomes, perceived difficulty and handling, perceived utility, and organizational aspects of the activity. Items were designed to explore both the theoretical understanding and the practical aspects of the activity. Specifically, they addressed students’ perceptions of their knowledge of tractography and white matter anatomy; their ability to navigate and interpret tractography data using TrackVis software; the perceived usefulness of the activity for their academic training; and organizational elements related to the structure and delivery of the practical sessions. The questionnaire is provided in Table 1.

**Table 1.** Learning Experience Questionnaire used to evaluate students’ perceptions of the virtual tractography activity.

|  | *DIFFICULTY AND HANDLING* | 1 | 2 | 3 | 4 | 5 |
| --- | --- | --- | --- | --- | --- | --- |
| D1 | The practical session was easy to follow |  |  |  |  |  |
| D2 | By the end of the practical session, I was able to use TrackVis independently |  |  |  |  |  |
| D3 | The workload (theory + practical session) was appropriate |  |  |  |  |  |
| D4 | During the practical session, I felt guided and able to follow the steps |  |  |  |  |  |
|  | *UTILITY* | 1 | 2 | 3 | 4 | 5 |
| U1 | The practical session helped me better understand neuroanatomy |  |  |  |  |  |
| U2 | Virtual dissection is a useful way to learn white matter anatomy |  |  |  |  |  |
| U3 | I would like more practical sessions like this in the course |  |  |  |  |  |
| U4 | I consider this practical session useful for my learning |  |  |  |  |  |
|  | *TEACHING ORGANIZATION* |  |  |  |  |  |
| O1 | The time allocated in class was sufficient |  |  |  |  |  |
| O2 | An additional practical session would have been useful |  |  |  |  |  |
| O3 | The instructor adequately addressed questions during the practical session |  |  |  |  |  |

No baseline assessment was conducted, as the questionnaire was designed to obtain post-activity feedback on students’ perceptions of the educational experience rather than to measure changes in knowledge. Given the highly specialized nature of tractography and the lack of prior formal instruction in this technique, a pre-intervention knowledge assessment was deemed uninformative. The questionnaire was administered after completion of the practical sessions. Responses to the Learning Experience Questionnaire were collected anonymously.

| EVALUATION QUESTIONNAIRE |  |  |  |  |  |  |
| --- | --- | --- | --- | --- | --- | --- |
| Instructions: Rate each statement from 1 to 5 according to your level of agreement<br>(1 = strongly disagree; 5 = strongly agree). |  |  |  |  |  |  |
|  | *LEARNING* | 1 | 2 | 3 | 4 | 5 |
| L1 | I understand what tractography is and what it represents |  |  |  |  |  |
| L2 | I know what a ROI is and how it is used to select fiber pathways |  |  |  |  |  |
| L3 | I have a better understanding of the anatomy and trajectory of the arcuate fasciculus after the practical session |  |  |  |  |  |
| L4 | I can interpret the results of the virtual dissection at a basic level |  |  |  |  |  |

### 2.5. Statistical analysis

All statistical analyses were conducted using RStudio (version 2025.09.2+418). For the analyses, items were grouped into four predefined dimensions reflecting different aspects of the educational experience: perceived learning outcomes (L1–L4), perceived difficulty and handling (D1–D4), perceived utility (U1–U4), and organizational aspects (O1–O3). The item assessing the need for an additional practical session (O2) was reverse-coded prior to analysis. Composite scores for each dimension were calculated as the mean of the corresponding items. Descriptive statistics (mean and standard deviation [SD]) were calculated for each dimension of the questionnaire. Internal consistency of each dimension was evaluated using Cronbach’s alpha. Pearson correlation analyses were conducted to examine the relationships between perceived learning and the other questionnaire dimensions.

To determine the relative contribution of each dimension to students’ perceived learning outcomes, a multiple linear regression model was conducted with perceived learning as the dependent variable and perceived difficulty and handling, perceived utility, and organizational aspects as predictors. Standardized regression coefficients were calculated to facilitate comparison of predictor importance. To further examine the relative contribution of each predictor, relative importance analyses were performed using the Lindeman, Merenda, and Gold (LMG) metric as implemented in *relaimpo* package (Grömping, 2006). Nested model comparisons were then conducted by sequentially removing each predictor from the full regression model and assessing changes in model fit using F-tests. Statistical significance was set at p < .05.

Regression assumptions were evaluated by inspecting diagnostic plots. Multicollinearity was assessed using variance inflation factors (VIF), residual normality using the Shapiro-Wilk test, and homoscedasticity using the Breusch-Pagan test. Because heteroscedasticity can bias standard errors, regression coefficients were additionally examined using heteroscedasticity-consistent (HC3) robust standard errors. Statistical significance was set at p < .05.

## 3. Results

### 3.1. Learning Experience Questionnaire results

Sixty-eight students completed the Learning Experience Questionnaire following participation in the virtual tractography activity. Students reported generally positive perceptions across the evaluated dimensions (Figure 3A) of the technology-enhanced learning experience. Perceived learning outcomes received the highest mean score (M = 3.44, SD = .85), followed by perceived utility (M = 3.35, SD = 1.05), perceived difficulty and handling (M = 3.33, SD = .88), and organizational aspects (M = 3.00, SD = .83).

**Figure 3.**
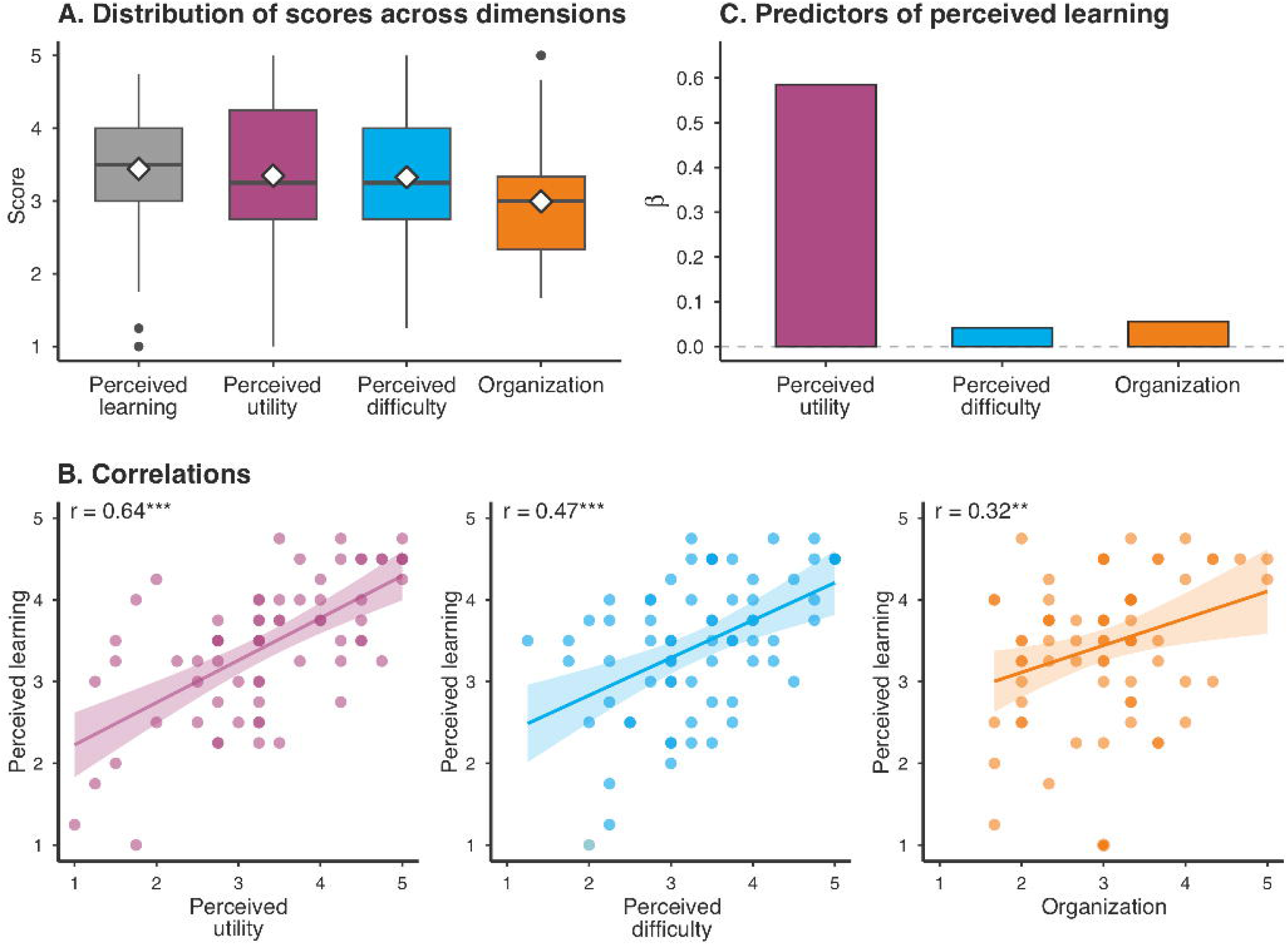
Results of students’ evaluation of the virtual tractography activity. **(A)** Distribution of mean scores across the dimensions assessed by the Learning Experience Questionnaire. Boxes represent the interquartile range, the horizontal black line indicates the median, and white diamonds denote the mean. **(B)** Scatterplots showing the relationships between perceived learning outcomes and perceived utility, perceived difficulty and handling, and organizational aspects. Solid lines represent fitted linear regression models, and shaded areas indicate 95% confidence intervals. Darker points represent overlapping observations. **(C)** Standardized regression coefficients (β) from the multiple regression model predicting perceived learning outcomes.

#### 3.1.1. Descriptive statistics and internal consistency

Internal consistency analyses indicated acceptable to good reliability across questionnaire dimensions. Cronbach’s alpha was 0.7 for perceived learning outcomes, 0.72 for perceived difficulty and handling, and 0.9 for perceived utility. Organizational aspects showed a lower but still acceptable level of internal consistency (α = .62), likely reflecting the heterogeneity of the items included in this dimension, which addressed different facets of the instructional experience, such as time allocation, the need for additional practice sessions, and instructor support.

#### 3.1.2. Correlations among dimensions

Pearson correlation analyses revealed significant positive correlations between perceived learning outcomes and all other dimensions of the learning experience (Figure 3B). Perceived utility showed the strongest correlation with perceived learning outcomes (r = .64, p < .001), followed by perceived difficulty and handling (r = .47, p < .001), and organizational aspects (r = .32, p < .007) (Figure 3B).

#### 3.1.3. Predictors of perceived learning

To determine the relative contribution of each educational dimension to students’ perceived learning outcomes, a multiple linear regression analysis was conducted with perceived learning as the dependent variable and perceived utility, perceived difficulty and handling, and organizational aspects as predictors. The model was statistically significant, explaining 41% of the variance in perceived learning outcomes (R² = .41, adjusted R² = .38, F (3,64) = 14.87, p < .001). Standardized regression coefficients indicated that perceived utility was the only significant predictor of perceived learning outcomes (β = .59, p = .001), whereas perceived difficulty and handling (β = .04, p = .78) and organizational aspects (β = .06, p = .65) did not contribute significantly to the model (Figure 3C). Relative importance analyses indicated that perceived utility accounted for 67.9% of the model’s explained variance, compared with 23.3% for perceived difficulty and handling, and 8.8% for organizational aspects. These percentages represent each predictor’s relative share of the total R², rather than percentages of total variance in perceived learning. Consistent with these findings, removing perceived utility from the model resulted in a significant reduction in model fit (F(1,64) = 20.16, p < .001), whereas the removal of perceived difficulty and handling (F(1,64) = .08, p = .78) or organizational aspects (F(1,64) = .21, p = .65) did not significantly affect explanatory power. Together, these findings identify perceived utility as the educational dimension most strongly associated with students’ perceived learning outcomes during the virtual tractography activity.

Assumption checks supported the adequacy of the regression model. There was no evidence of problematic multicollinearity (all VIFs < 2.5), and residuals were approximately normally distributed (Shapiro-Wilk p = .41). Although mild heteroscedasticity was detected (Breusch-Pagan p = .016), the application of HC3 robust standard errors did not alter the magnitude or statistical significance of any regression coefficient. In particular, perceived utility remained the only significant predictor of perceived learning outcomes (p < .001), whereas perceived difficulty and handling, and organizational aspects remained non-significant. These findings indicate that the observed pattern of results was robust to potential violations of homoscedasticity assumptions.

## 4. Discussion

The present study evaluated a virtual tractography-based neuroanatomy activity implemented within a Psychobiology course for undergraduate Psychology students. Overall, students reported moderately positive perceptions across the evaluated dimensions, with perceived learning outcomes receiving the highest mean ratings. More importantly, perceived utility emerged as both the strongest correlate and the only significant predictor of perceived learning outcomes. Relative importance analyses further showed that perceived utility accounted for the largest proportion of explained variance. These findings indicate that perceived learning was most strongly associated with the extent to which students considered the activity useful. In contrast, perceived difficulty and organizational aspects did not make significant independent contributions to the model. From a practical standpoint, although tractography is a technically demanding neuroimaging technique with which students had no prior experience, the generally positive ratings suggest that research-oriented neuroimaging tools can be successfully integrated into undergraduate neuroanatomy teaching.

The significant role of perceived utility observed in the present study fits well with educational theories that emphasize the importance of task value in learning (see Eccles & Wigfield, 2002, for a review). In Expectancy-Value Theory (Eccles et al., 1983; Wigfield & Eccles, 2000), subjective task value refers to the value and importance that students attribute to an educational activity and comprises four components: attainment value, intrinsic value, utility value, and cost. The finding that students who perceived the tractography activity as more useful reported greater perceived learning is consistent with the broader role attributed to utility value in Expectancy-Value Theory, defined as the perceived usefulness of a task for achieving current or future goals, which constitutes an important motivational resource that can support learning and academic engagement.

A complementary perspective is provided by the Four-Phase Model of Interest Development (Hidi & Ann Renninger, 2006; Hidi & Renninger, 2026). Within this framework, interest development begins with situational interest, a psychological state elicited by features of a task or learning environment, and may subsequently develop into individual interest, a more stable predisposition toward a domain. Situational interest is initially triggered by salient features such as novelty, personal relevance, intensity, or surprising information, plausibly reflected in the novelty of 3D tractography visualization and the interactivity of the software (Guo & Fryer, 2025). However, for learning benefits to be sustained, situational interest must be maintained through activities perceived as meaningful and relevant. In this regard, although novelty and interactivity may have contributed to students’ initial engagement, these factors were not directly assessed in the present study. Our findings instead indicate that perceived utility was the dimension most strongly associated with perceived learning. Although the technological characteristics of the activity may have initially captured students’ attention, perceived utility emerged as the strongest predictor of perceived learning outcomes, indicating that students benefited most when they recognized the relevance of the activity for understanding neuroanatomy. Although the present study did not assess longer-term changes in students’ interest, this interpretation is consistent with the proposition that repeated learning experiences that successfully elicit and maintain situational interest may support the gradual development of more enduring forms of interest in a domain (Guo & Fryer, 2025; Hidi & Ann Renninger, 2006)

The present findings may be particularly relevant within Psychology education. Neurophobia has been well documented among medical trainees (Flanagan et al., 2007; McCarron et al., 2014), but research in non-medical disciplines remains comparatively limited. Recent evidence, however, indicates that Psychology students also experience difficulties and apprehension when learning neuroanatomy, suggesting that neurophobia may extend beyond medical education (Garcés-Arilla et al., 2025; Garces-Arilla et al., 2026). Unlike medical students, Psychology students may not readily perceive the relevance of neuroanatomical knowledge to their future professional practice (Garcés-Arilla et al., 2025). This may further compound the challenges posed by the inherent spatial and structural complexity of neuroanatomy (Mendez-Lopez et al., 2022). Importantly, evidence suggests that neurophobia is more strongly associated with modifiable educational factors than with the inherent complexity of neuroscience itself (Garces-Arilla et al., 2026). Educational strategies aimed at increasing interest, including computer-assisted learning resources, have likewise been associated with reduced neurophobia and improved neuroanatomy learning (Javaid et al., 2018). Our findings identify perceived utility as a potentially relevant educational dimension that warrants further investigation. By introducing students to a contemporary, hands-on neuroimaging technique actively used in cognitive and clinical neuroscience research (Forkel et al., 2022; Johansen-Berg & Behrens, 2006), the tractography activity may have strengthened students’ perception that neuroanatomical knowledge connects to authentic scientific and clinical research and practice. This interpretation is consistent with evidence that utility-value interventions can increase interest, engagement, achievement, and persistence (Hulleman et al., 2010; Hulleman & Harackiewicz, 2009). From a practical perspective, these findings suggest that future studies should examine whether explicitly communicating the purpose and potential educational value of neuroanatomical activities can influence students’ perceived utility and, in turn, their learning experiences.

Beyond its potential to increase the perceived relevance of neuroanatomy, virtual tractography-based dissection may be particularly well suited as a hands-on educational complement for teaching the complex spatial organization of white matter pathways. White matter anatomy is particularly challenging for students because fiber pathways form a highly complex network of long-range connections with overlapping trajectories, crossing fibers, and intricate spatial relationships that are difficult to appreciate using static two-dimensional representations. Interactive visualization tools have been shown to facilitate neuroanatomy learning, increase engagement, and be preferred over traditional two-dimensional methods (Elsayed et al., 2025; Mendez-Lopez et al., 2022), while also having the potential to trigger situational interest (Guo & Fryer, 2025). By allowing students to manipulate tractography data, define ROIs, reconstruct fiber pathways, and explore them from multiple viewpoints, the present activity may support the construction of more coherent mental representations of white matter organization than conventional static images alone.

Several limitations should be acknowledged. First, the study was conducted in a single cohort of first-year undergraduate Psychology students at a single university, which may limit the generalizability of the findings to other educational contexts. Second, the evaluation was based primarily on students’ self-reported perceptions and did not include a comprehensive pre-post assessment of neuroanatomical knowledge. Consequently, it is not possible to determine whether perceived learning accurately reflected objective learning gains. Third, the practical activity focused exclusively on the arcuate fasciculus and therefore may not fully represent the educational potential of tractography-based instruction across different white matter pathways and neuroanatomical systems. Finally, it was not possible to examine whether the strong relationship between perceived utility and perceived learning outcomes extended to objective indicators of learning or academic performance. Finally, because questionnaire responses were collected anonymously, it was not possible to examine whether the observed relationship between perceived utility and perceived learning extended to objective indicators of academic performance. Future studies should determine whether tractography-based learning activities improve objective neuroanatomical knowledge and whether the strong association observed between perceived utility and perceived learning is also reflected in academic achievement. Longitudinal research could further examine whether repeated exposure to technology-enhanced neuroanatomy activities influences students’ interest in neuroscience and neurophobic attitudes over time.

## 5. Conclusions

Virtual tractography appears to be a feasible and well-accepted strategy for introducing complex white matter anatomy in undergraduate Psychology education. Students reported generally positive perceptions of the learning experience, and perceived utility, rather than difficulty and handling or organization of the activity, emerged as the dimension most strongly associated with perceived learning outcomes and the only significant predictor. These findings suggest that helping students recognize the relevance and usefulness of neuroanatomical content, for instance by explicitly linking it to research and clinical practice, may be an important consideration when designing neuroscience learning experiences and addressing some of the challenges underlying neurophobia in Psychology students. The activity also illustrates how research-oriented neuroimaging tools can be incorporated into undergraduate teaching when accompanied by structured instruction, demonstrations, and guided practice, even among students with no previous experience with such technologies. Beyond neuroanatomy, the findings highlight the potential of technology-enhanced learning experiences that combine interactive visualization with authentic scientific tools to support the learning of complex spatial concepts in undergraduate STEM education. The capacity of tractography to provide direct three-dimensional visualization and hands-on reconstruction of white matter pathways may therefore offer educational advantages that extend beyond traditional anatomy instruction.

## 6. Declarations

## Availability of data and materials

The video tutorial and tractography dissection guide used in the practical activity are publicly available in Zenodo repository: https://doi.org/10.5281/zenodo.22129828. The datasets used and/or analysed during the current study are available from the corresponding author on reasonable request.

## Author contributions

D.L-B: conceptualization, formal analysis, investigation, funding acquisition, visualization, Writing -Original Draft Preparation, Writing – Review & Editing. MJT-P: conceptualization, formal analysis, Writing – Review & Editing.

## Acknowledgements

Not aplicable.

## Footnotes

1 Traditional plastic brain models may sometimes promote simplified spatial representations. For example, students frequently refer to the insular cortex as “the lid” because many plastic brain models depict it as a removable cortical piece that exposes the basal ganglia underneath. While pedagogically useful, such representations may obscure the fact that the insula is not an independent structure but rather a continuous region of cerebral cortex located beneath the opercula. Interactive three-dimensional neuroimaging tools may help students build more accurate mental representations of these complex anatomical relationships.

