## Supplementary material for "Technology-enhanced learning in undergraduate neuroscience education: tractography-based virtual dissection in psychology": Figure S1

**Teaching neuroanatomy through tractography-based virtual dissection:**

**a learning experience in undergraduate Psychology education**

**Figure S1.**

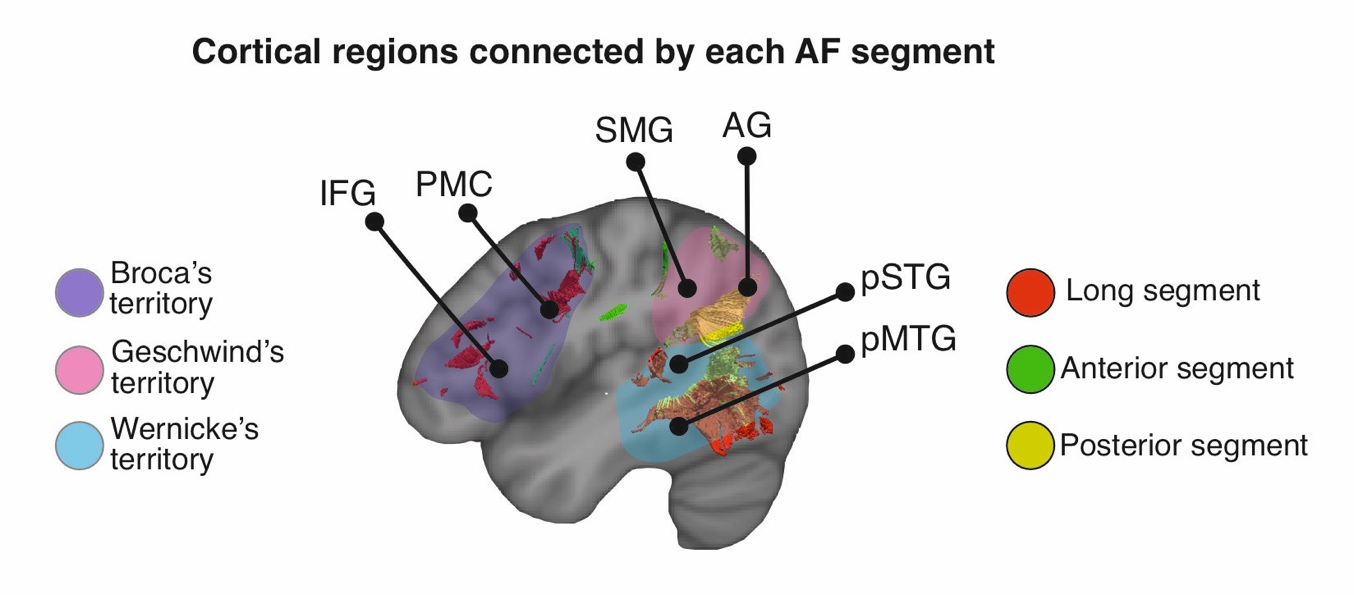

**Figure caption**

**Figure S1.** Cortical regions associated with the three segments of the arcuate fasciculus. Schematic representation of the cortical territories connected by the long (red), anterior (green), and posterior (yellow) segments of the arcuate fasciculus. Colored cortical regions indicate Broca’s territory (purple), Geschwind’s territory (pink), and Wernicke’s territory (blue). IFG = inferior frontal gyrus; PMC = premotor cortex; SMG = supramarginal gyrus; AG = angular gyrus; pSTG = posterior superior temporal gyrus; pMTG = posterior middle temporal gyrus.
